# Restoring Cholesterol Efflux in Vascular Smooth Muscle Cells Transitioning into Foam Cells Through Liver X Receptor Activation

**DOI:** 10.1101/2025.01.22.634420

**Authors:** Carla Borràs, Noemí Rotllan, Raquel Griñán, David Santos, Marta Mourín, Begoña Soto, Mercedes Camacho, Mireia Tondo, Marina Canyelles, Francisco Blanco-Vaca, Joan Carles Escolà-Gil

## Abstract

**Objective:** Macrophage foam cells derived from vascular smooth muscle cells (VSMCs) account for 30–70% of foam cells in atherosclerotic lesions. Liver X receptor (LXR) agonists promote high-density lipoprotein (HDL)-mediated cholesterol efflux from macrophages. This study aimed to investigate the effects of LXR activation on the reverse cholesterol transport (RCT) rate from VSMCs to feces *in vivo*.

**Approach and Results:** Cholesterol efflux induced by serum and HDL was evaluated in human and mouse VSMCs treated with the LXR agonist T090137 before and after methyl-β-cyclodextrin (MBD)-cholesterol loading. Additional experiments included treatment with an acyl-coenzyme A: cholesterol acyltransferase (ACAT) inhibitor. Cholesterol-radiolabeled VSMCs were injected into the peritoneal cavity of mice, and RCT was assessed by measuring radiolabeled cholesterol in serum, liver, and feces over 48 hours. Serum and HDL induced cholesterol efflux at similar levels in both human and mouse VSMCs. Cholesterol efflux was significantly reduced following MBD-cholesterol loading; however, treatment with the LXR agonist significantly enhanced efflux. Radiolabeled foam-like VSMCs injected into mice exhibited impaired cholesterol transfer to serum, HDL, and feces compared to non-lipid-laden VSMCs. Pre-treatment with the LXR agonist increased radiolabeled cholesterol levels in serum and HDL and doubled its fecal excretion. Additionally, LXR activation restored RCT from MBD-cholesterol-loaded VSMCs to feces, reaching levels comparable to those of non-lipid-laden cells. Treatment with an ACAT inhibitor fully restored RCT rates in foam-like VSMCs, and the combination of the ACAT inhibitor and the LXR agonist further enhanced RCT.

**Conclusions:** HDL-mediated cholesterol efflux is significantly impaired in VSMCs during their transition into foam cells. Pharmacological activation of LXR enhances RCT from VSMCs to feces *in vivo* and restores the impaired RCT from transitioning VSMCs. The combination of LXR agonists and ACAT inhibitors holds promise as a synergistic therapeutic approach to restoring cholesterol homeostasis in lipid-laden VSMCs, offering potential strategies to mitigate atherosclerosis.

**Highlights:**

- LXR activation enhances cholesterol efflux in VSMCs *in vitro*, even after their transition into foam cells.
- VSMCs transitioning into foam cells exhibit reduced cholesterol transfer to HDL and feces in mice.
- LXR agonist treatment enhances reverse cholesterol transport (RCT) from VSMCs to feces *in vivo*.
- Selective ACAT inhibition restores RCT in foam-like VSMCs, with further enhancement observed upon LXR activation.

## Introduction

Atherosclerosis, the leading cause of cardiovascular disease, is characterized by the progressive deposition and accumulation of cholesterol and fibrous material within the tunica intima of arterial walls.^1^ Vascular smooth muscle cells (VSMCs) are estimated to constitute more than 90% of all cells in human atherosclerotic plaques and are also present in the pre-atherosclerotic arterial intima. The onset of atherosclerosis in humans is preceded by the formation of a thick intimal layer, known as diffuse intimal thickening, which is composed of VSMCs and their secreted proteoglycans.^2^ Despite this, most descriptions of developing and advanced atherosclerosis in major reviews of the disease pathogenesis emphasize the role of macrophages, often portraying VSMCs as playing a relatively minor role. In these accounts, the contribution of VSMCs is typically confined to their involvement in forming the fibrous cap over lesions.^3^

Foam cells are recognized as the primary contributors to atheroma plaque formation, with their origins traditionally attributed to macrophages. However, studies have shown that at least 50% of foam cells in human coronary artery atheromas and 30–70% in mouse atheromas are derived from VSMCs.^4,5^ This phenomenon is partly explained by the loss of VSMC markers, such as α-smooth muscle actin and myosin heavy chain, coupled with an increased expression of macrophage markers, including CD68 and Mac-2. These changes have led to the underestimation of VSMC-derived foam cells in previous analyses.^6^

High-density lipoproteins (HDL) play a pivotal role in reverse cholesterol transport (RCT) by facilitating the efflux of excess cholesterol from peripheral tissues to the liver for excretion via bile.^7^ While much attention in the early steps of RCT has focused on elucidating cholesterol efflux mechanisms in macrophages, investigations have also extended to VSMCs. Cholesterol efflux, the initial step of the RCT pathway, is mediated by various cholesterol transporters, with ATP-binding cassette subfamily A member 1 (ABCA1) playing a predominant role in VSMCs. ABCA1 expression and its interaction with apolipoprotein (APO) A1 are significantly reduced in human VSMCs, particularly in advanced atherosclerosis compared to early-stage disease.^4,8^ A similar reduction is observed in intimal-like VSMCs relative to those from the medial layer.^9^ Consistent with these findings, foam cells derived from VSMCs in APOE-deficient mice exhibit diminished ABCA1 expression and impaired cholesterol efflux to APOA1 compared with foam cells of leukocyte origin.^10^ In addition, VSMC-derived foam cells demonstrate greater impairment in autophagy, a cellular process involved in intracellular cholesterol trafficking to the efflux pathway, when compared to macrophage-derived foam cells.^11^ Cholesterol efflux promoted by HDL or APOA1 has been shown to reverse phenotypic changes induced by cholesterol loading in VSMCs, further underscoring its therapeutic potential.^12^

Liver X Receptors (LXRs) are oxysterol-activated transcription factors that regulate a network of genes to promote the coordinated mobilization of excess cholesterol from cells and the body.^13^ Treatment with LXR agonists has been shown to enhance cholesterol efflux from macrophages, partly through the upregulation of ABCA1 expression.^14^ The LXR agonist T0901317 has further demonstrated the ability to promote RCT from macrophages to feces in mice by increasing plasma cholesterol efflux capacity.^15^ Macrophage LXR activity is a crucial, albeit not exclusive, component of the entire RCT pathway in response to LXR agonists, as evidenced by studies in mice.^16^ Bone marrow transplantation experiments have also confirmed that LXR activation in macrophages is essential for the atheroprotective effects of T0901317 in murine models.^17^ However, despite the induction of ABCA1 expression by LXR agonists, APOA1-mediated cholesterol efflux could not be restored in cultured epithelioid VSMCs derived from rat thoracic aortas.^9^

In contrast to macrophages, no *in vivo* studies have evaluated RCT from VSMCs transitioning into foam cells or whether this process can be reversed by LXR activation, as well as the molecular mechanisms underlying these alterations. In this study, we evaluated the capacity of serum and HDL to enhance cholesterol efflux from cultured human and murine VSMCs, both prior to and following their transformation into foam cells via methyl-β-cyclodextrin (MBD)-cholesterol loading. Additionally, we directly tested the hypothesis that LXR activation enhances the RCT rate from VSMCs to feces *in vivo*, while also investigating the role of acyl-coenzyme A: cholesterol acyltransferase (ACAT) inhibition in restoring this pathway.

## Methods

### Cell culture

Primary VSMCs were isolated from human aortas using the explant technique. Aortic tissues were obtained from multi-organ donors, with informed consent obtained prior to tissue collection and delivery. Briefly, aortic tissues were thoroughly rinsed in phosphate buffered saline (PBS, Sigma Aldrich/Merck, St Louis, MO) containing antibiotics to remove contaminating blood cells. Subsequently, the adipose tissue was carefully excised, and the medial layer was dissected from the adventitia and cut longitudinally. The endothelium was removed by scraping with forceps, and the medial layer was cut into 1-2 mm^3^ pieces before being transferred to culture plates, which were incubated at 37ªC for 2 hours to allow cell adhesion. Once the tissue attached, 5 mL of Dulbecco’s Modified Eagle Medium (DMEM) high glucose with L-glutamine and sodium pyruvate (Corning, Corning, NY) supplemented with 10% fetal bovine serum (FBS; Pan Biotech, Aidenbach, Germany) and 100 U/mL penicillin/streptomycin (Corning, Corning, NY) were carefully added. Cells were incubated at 37°C in a humidified atmosphere containing 5% CO_2_ for 1-2 weeks and the culture medium was replaced every 48 hours. When cells reached confluence, the medial tissue pieces were removed, and the cells were trypsinized and passaged according to standard protocols. All VSMCs used in this study were at passage 4-8.

Due to the large quantity of VSMCs required for *in vivo* experiments, immortalized mouse aortic VSMCs (ATCC^®^ CRL-2797™, American Type Culture Collection, Manassas, VA) were cultured in DMEM supplemented with 10% FBS and 0.2 g/L G-418 (InvivoGen, San Diego, CA). The cells were maintained in 75 cm^2^ cell culture flasks and incubated at 37°C in a humidified atmosphere containing 5% CO_2_. The culture medium was replaced every 48 hours, and cells were trypsinized when they reached confluence.

For the experiments, human or mouse VSMCs were seeded and cultured in complete medium for 24 to 72 hours. Subsequently, the cells were exposed to 20 μg/mL MBD-cholesterol (Sigma-Aldrich/Merck, St. Louis, MO) in 5% FBS-containing medium for 48 hours to induce a foam cell phenotype, representing the VSMC cholesterol-loading phase. Non-loading control cells were incubated with the same medium without MBD-cholesterol. Afterwards, the VSMCs underwent an LXR activation step, which involved treatment with an LXR agonist T0901317 (Cayman Chemicals, Ann Arbor, MI) at a concentration of 2μmol/L, dissolved in dimethyl sulfoxide (Sigma-Aldrich/Merck, St. Louis, MO), for 18 hours. Untreated control cells were incubated with dimethyl sulfoxide alone as the vehicle (0.1% v/v). Non-loading and untreated control cells were considered the baseline conditions for the experiments.

For those experiments involving ACAT inhibition, the ACAT inhibitor Sandoz 58-035 (Santa Cruz Biotechnology, Dallas, TX) was added to the culture medium at a concentration of 10 μmol/L during all experimental steps.

### Mice and diet

Wild-type C57BL/6J mice were obtained from Jackson Laboratories (Bar Harbor, ME; #000664) and housed in a temperature-controlled environment (22 °C) with a 12-hour light/dark cycle. Food and water were provided ad libitum. The sample size for each group was determined based on statistical parameters (α=0.05, power=80%) and an estimated effect size of 5,000 counts per minute in the fecal excretion of VSMC-derived cholesterol. To minimize bias, mice with the same age were randomized into groups of equal size, and all animals were included in the analyses. Experiments were conducted in a blinded manner with respect to the origin of the specimens. No mice were excluded in the study.

The experimental procedures were reviewed and approved by the Institutional Animal Care and Use Committee of the Sant Pau Research Institute and authorized by the Animal Experimental Committee of the local government authority (Generalitat de Catalunya, authorization No. 10626) in accordance with the Spanish Law (RD 53/2013) and European Directive 2010/63/EU. Procedures were conducted at the Animal Experimentation Service, ISO 9001:2015 certified.

### Oil Red O staining

Oil Red O staining was performed to visualize the accumulation of neutral lipids within VSMCs. Cells were seeded into 6-well plates, cultured in complete medium for 24 hours, and treated under previously described experimental conditions. Following treatment, the cell medium was removed, and the cells were washed twice with PBS before fixation with 4% paraformaldehyde (Thermo Fisher Scientific, Waltham, MA) for 10 minutes. After fixation, paraformaldehyde was removed, and the cells were washed twice with PBS. Cells were stained with an Oil Red O working solution prepared by mixing 4 parts distilled water with 6 parts of a 5 g/L Oil Red O stock solution (Sigma-Aldrich/Merck, St. Louis, MO) dissolved in isopropanol (Sigma-Aldrich/Merck, St. Louis, MO). The staining process was carried out for 1 hour. After staining, cells were washed twice with distilled water, counterstained with a 1 g/L hematoxylin solution (Sigma-Aldrich/Merck, St. Louis, MO) for 1 minute, and rinsed again with distilled water. Images of the cells were captured using an inverted fluorescence phase-contrast microscope (Leica Microsystems, Wetzlar, Germany).

### Quantitative Real Time-PCR analysis

For quantitative real time PCR analyses, total cellular RNA was extracted using the EZ-10 DNAaway RNA Miniprep kit (Bio Basic Inc., Markham, Ontario, Canada) and quantified by NanoDrop-2000 spectrophotometry (Thermo Fisher Scientific, Waltham, MA). cDNA was synthesized using EasyScript First-Strand cDNA Synthesis SuperMix (Transgen Biotech, Beijing, China). Quantitative Real-Time PCR amplification was performed using the GoTaq® Probe qPCR Master Mix (Promega, Madison, WI) with specific TaqMan® probes (Applied Biosystems, Foster City, CA). Probes for human genes were *ABCA1* (*Hs01059118_m1*), *CD68* (*Hs00154355_m1*), *Transgrelin (TAGLN*, *Hs01038780_m1*) and *Glyceraldehyde-3-phosphate Dehydrogenase* (*GAPDH*, *Hs99999905_m1*), and for mouse genes were *Abca1* (*Mm00442646_m1*), *Cd68* (*Mm03047343_m1*), *Tagln* (*Mm00441661_g1*) and *Actin* (*Mm00607939_s1*).

Reactions were conducted on a CFX96^TM^ Real-Time System (Bio-Rad, Hercules, CA) following the manufacturer’s instructions. Thermal cycling conditions consisted of an initial denaturation step at 95 °C for 10 minutes, followed by 40 cycles of 95 °C for 15 seconds and at 65 °C for 1 minute. Relative mRNA expression levels were calculated using the ΔΔCt method, with human gene expression normalized to *GAPDH* and mouse gene expression normalized to *Actin*.

### Cholesterol efflux

*In vitro* cellular cholesterol efflux was evaluated using an adapted radiochemical method.^18^ Human primary cultured VSMCs or immortalized mouse aortic VSMC were seeded in 6-well plates and cultured in complete medium for 24 hours. VSMCs were then labeled with DMEM containing 1 µCi/well of [1α,2α(n)-^3^H]cholesterol (Perkin Elmer, Boston, MA) and 5% FBS under baseline conditions or during the cholesterol-loading phase for 48 hours. After labeling, cells were equilibrated overnight in medium containing 0.2% free fatty acid bovine serum albumin (Sigma Aldrich/Merck, St Louis, MO). Optionally, LXR activation was performed during this step. The next day, VSMCs were extensively washed, and medium containing cholesterol acceptors was added to assess cholesterol efflux over 4 hours.

Cholesterol acceptors included pooled serum samples from normolipemic individuals (human acceptors) or wild-type C57BL/6J mice (mouse acceptors). For the experiments, 25 μL of serum (2.5%, v/v), 75 μL of serum after precipitation of APOB-containing lipoproteins using 0.44 mmol/L phosphotungstic acid and 20 mmol/L magnesium chloride (Merck, Darmstadt, Germany), or HDL (25 µg/mL of APOA1) isolated by ultracentrifugation (density range 1.063–1.210 g/mL) were used.^18^ Human plasma samples from healthy donors were obtained following the standards for medical research involving human subjects as outlined in the Declaration of Helsinki. The study protocol was approved by the Ethical Committee of Hospital de la Santa Creu i Sant Pau (protocol code IIBS-APO-2013-105).

The percentage of cholesterol efflux was calculated by dividing the radiolabeled cholesterol present in the medium by the total radiolabeled cholesterol (the sum of cholesterol in the medium and cell-associated cholesterol). Liquid scintillation counting was used to quantify radiolabeled cholesterol at the conclusion of the experiments.

### *In vivo* VSMC-to feces RCT assay

Eight- to ten-week-old wild-type male and female mice were fed a Western-type diet containing 21% fat and 0.2% cholesterol (TD.88137, Harlan Teklad, Madison, WI) for 4 weeks. To activate LXR systemically, mice were administered an intragastric dose of the synthetic LXR agonist T0901317 (10 mg/kg in 1% w/v carboxymethylcellulose, Sigma-Aldrich/Merck, St. Louis, MO) during the final 12 days of the diet, including the 48 hours of the RCT assay. Control animals received a vehicle (1% w/v carboxymethylcellulose).

Immortalized mouse aortic VSMCs were cultured in 75 cm^2^ cell culture flasks in complete medium for 72 hours. Subsequently, the cells were incubated with 5 µCi/mL of [1α,2α(n)-³H]cholesterol and 5% FBS under baseline conditions or during the cholesterol-loading phase for 48 hours. For experiments involving ACAT inhibition, the ACAT inhibitor was added to the culture medium during the cholesterol-loading phase. Following incubation, the cells were washed and equilibrated overnight in medium containing 0.2% free fatty acid bovine serum albumin. Optionally, LXR activation was performed during the equilibration step. The radiolabeled mouse VSMCs were trypsinized, resuspended in PBS and intraperitoneally injected into the mice (average of 1.3·10^6^ VSMCs containing 1.0·10^6^ cpm per mouse; cell viability was ≥92%, as measured by trypan blue staining).

Forty-eight hours before the end of the experiment, mice were randomized into four groups, with three types of experimental approaches performed:

First Experimental Approach:

Group 1: Mice injected with radiolabeled VSMCs under baseline conditions.

Group 2: Mice injected with radiolabeled VSMCs under cholesterol-loading conditions.

Group 3: Mice injected with radiolabeled VSMCs pre-treated with the LXR agonist.

Group 4: Mice injected with radiolabeled VSMCs under cholesterol-loading conditions and pre-treated with the LXR agonist.

Second Experimental Approach:

Group 1: Mice treated with the vehicle solution and injected with radiolabeled VSMCs under baseline conditions.

Group 2: Mice treated with the vehicle solution and injected with radiolabeled VSMCs under cholesterol-loading conditions.

Group 3: Mice treated with the LXR agonist and injected with radiolabeled VSMCs under baseline conditions.

Group 4: Mice treated with the LXR agonist and injected with radiolabeled VSMCs under cholesterol-loading conditions.

Third Experimental Approach:

Group 1: Mice injected with radiolabeled VSMCs under baseline conditions.

Group 2: Mice injected with radiolabeled VSMCs under cholesterol-loading conditions.

Group 3: Mice injected with radiolabeled VSMCs under cholesterol-loading conditions and pre-treated with the ACAT inhibitor.

Group 4: Mice injected with radiolabeled VSMCs under cholesterol-loading conditions and pre-treated with both the ACAT inhibitor and the LXR agonist.

Following the injections, mice were individually housed, and stools were collected over the subsequent 48 hours. At the end of this period, the mice were weighted, and their food intake was recorded. Mice were then euthanized via exsanguination through cardiac puncture, and their livers were harvested for further analysis. Serum [^3^H]cholesterol radioactivity was measured by liquid scintillation counting. HDL [^3^H]cholesterol levels in serum were determined after precipitating APOB-containing lipoproteins using 0.44 mmol/L phosphotungstic acid and 20 mmol/L magnesium chloride. Hepatic and fecal lipids were extracted using isopropyl alcohol-hexane (2:3 v/v, Sigma Aldrich/Merck, St Louis, MO) as previously described.^19^ The lipid layers were collected, and [^3^H]cholesterol was quantified via liquid scintillation counting. The [^3^H]tracer detected in the fecal bile acids was combined with the [³H]cholesterol detected in feces to determine total fecal [³H]tracer. The amount of [^3^H]tracer was expressed as a fraction of the injected dose.

### Thin-layer chromatography analysis

Thin-layer chromatography analysis was performed to separate [³H]unesterified cholesterol from [³H]cholesteryl esters from immortalized mouse aortic VSMCs labeled with 1 µCi/well of [1α,2α(n)-3H]cholesterol (Perkin Elmer, Boston, MA) under baseline or cholesterol-loading conditions, with or without the LXR agonist T0901317 and the ACAT inhibitor Sandoz 58-035 treatment as previously described.

Cellular lipids were extracted by adding 1 mL of hexane-isopropanol (3:2 v/v, Sigma-Aldrich/Merck, St. Louis, MO) to the cells and incubating overnight at 4°C. The following day, the extract was collected, and 300 μL of 0.5 mol/L sodium sulfate (Sigma-Aldrich/Merck, St. Louis, MO) were added. The mixture was gently mixed for 10 minutes and then centrifuged at 3000 rpm for 10 minutes at 4°C, to separate the hexane and the isopropanol in two layers. The upper hexane layer containing lipids was collected and evaporated at room temperature. The dried lipid pellet was resuspended in 25 μL of chloroform (Sigma-Aldrich/Merck, St. Louis, MO) and applied onto a thin-layer chromatography plate (Sigma-Aldrich/Merck, St. Louis, MO). After, the plate was placed into a glass tank containing the mobile phase solution composed of hexane-diethyl ether-ethyl acetate (50:50:1.5, v:v:v, Sigma-Aldrich/Merck, St. Louis, MO) until the solvent front reached 5 cm from the top of the plate. Next, the plate was left overnight at room temperature and subsequently placed in a tank with iodine crystals (Sigma-Aldrich/Merck, St. Louis, MO) for 15 minutes to stain the lipids. The iodine-stained lipid bands were scraped from the plate and quantified by liquid scintillation counting.

### Lipid parameters analyses

Total serum cholesterol and triglycerides were quantified enzymatically using commercial assays, adapted for the COBAS 6000/501c autoanalyzer (Roche Diagnostics, Rotkreuz, Switzerland). HDL cholesterol levels in serum were determined after precipitating APOB-containing lipoproteins, as described above. A liver sample was also collected to evaluate hepatic lipid content. Liver lipids were extracted using a mixture of isopropyl alcohol and hexane (2:3, v/v). The lipid layer was then collected, evaporated, and resuspended in 0.5% (w/v) sodium cholate (Serva, Heidelberg, Germany). Cholesterol and triglyceride concentrations were measured using commercial kits adapted for the COBAS 6000/501c autoanalyzer.

### Statistical methods

D’Agostino & Pearson, Shapiro-Wilk, and Kolmogorov-Smirnov normality tests were performed to assess Gaussian distribution. The Student’s t-test was used to compare the differences between two groups. GraphPad Prism version 8.0.2 for Windows (GraphPad Software, San Diego, CA) was used to perform all statistical analyses. A p-value ≤ 0.05 was considered statistically significant.

### Data availability

The data, analytical methods, and study materials will be available to other researchers for purposes of reproducing the results or replicating the procedure upon reasonable request. Source data are provided with this paper.

## Results

### LXR agonist enhances cholesterol efflux in VSMCs and foam-like cells *in vitro* despite foam cell-induced reduction

We first confirmed that both primary VSMCs, derived from human aortas using the explant technique, and immortalized mouse aortic VSMCs exposed to MBD-cholesterol effectively mimicked the key characteristics of VSMCs transformed into foam-like cells (see the experimental outline in **Figure 1A**). As expected, VSMC cholesterol-loading increased lipid droplet accumulation in both human and mouse VSMCs, as evidenced by Oil Red O staining (**Figure 1B-C**). Furthermore, MBD-cholesterol treatment upregulated the mRNA expression of the macrophage marker *CD68*, while reducing the expression of the VSMC canonical marker *TAGLN* in human primary VSMCs (**Figure 1D-E**). Similarly, the expression of *Cd68* and *Tagln* followed the same pattern in mouse VSMCs when exposed to MBD-cholesterol, independently of LXR activation (**Figure F-G**).

**Figure 1:**
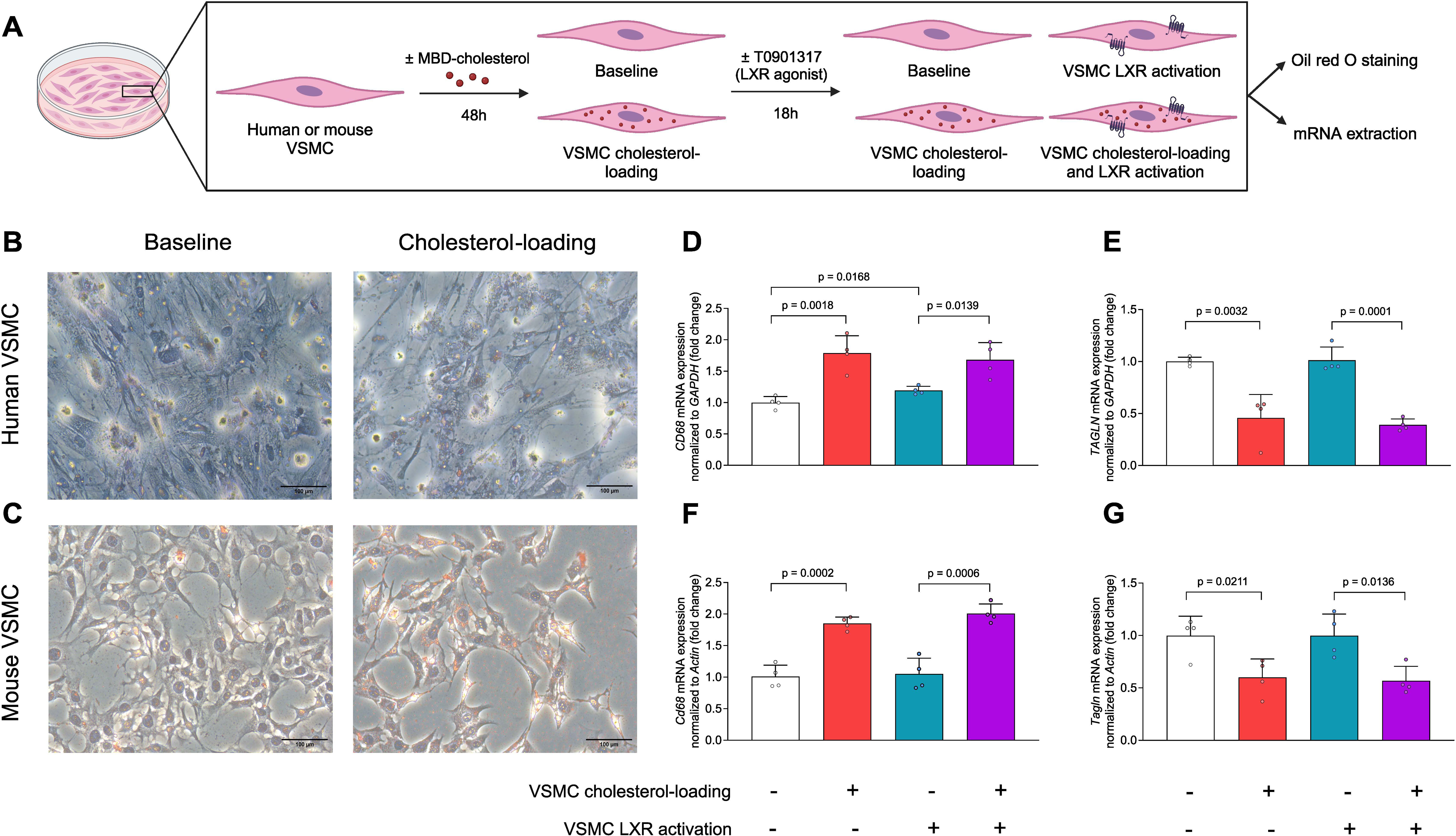
Human and Mouse VSMCs Exposed to MBD-Cholesterol Effectively Mimic Foam Cell Transformation Characteristics. (A) Primary VSMCs derived from human aortas, and immortalized mouse aortic VSMCs were cultured under baseline conditions or exposed to 20 μg/mL MBD-cholesterol, representing the VSMC cholesterol-loading phase, for 48 hours. VSMCs were then treated with either the vehicle or 2 µmol/L of the LXR agonist T0901317 for 18 hours, and subsequently Oil Red O staining or mRNA extraction was performed. (B) Oil Red O staining of human VSMCs under baseline or VSMC cholesterol-loading conditions. (C) Oil Red O staining of mouse VSMCs under baseline or VSMC cholesterol-loading conditions. (D, E) Human VSMCs, under baseline or VSMC cholesterol-loading conditions, were treated with either the vehicle or 2 µmol/L of the LXR agonist T0901317 for 18 hours. mRNA was isolated, and quantitative real-time PCR was performed to measure the relative mRNA expression of *CD68* and *TAGLN*. (F, G) mRNA was isolated from mouse VSMCs, and quantitative real-time PCR was performed to measure the relative mRNA expression of *Cd68* and *Tagln*. Baseline expression for each cell type was set to 1 arbitrary unit and subsequent expression levels were expressed as fold changes. Human gene expression was normalized to *GAPDH* and mouse gene expression was normalized to *Actin*. Data are presented as the mean ± SD from 4 independent experiments per group. Unpaired t-tests were used to compare mRNA expression between groups. GAPDH: Glyceraldehyde-3-phosphate Dehydrogenase, LXR: Liver X Receptor, MBD: methyl-β-cyclodextrin, TAGLN: Transgelin, VSMC: vascular smooth muscle cell.

We evaluated the potential of serum acceptors to promote cholesterol efflux in both human and mouse VSMCs, as outlined in the experimental design depicted in **Figure 2A**. Serum and APOB-depleted serum (containing HDL, preβ-HDL, remodeling factors, and albumin) similarly promoted cholesterol efflux in human VSMCs (**Figures 2B–C**). Cholesterol efflux was diminished when human VSMCs were exposed to MBD-cholesterol. Importantly, treatment with an LXR agonist enhanced cholesterol efflux in human VSMCs, including those that had transitioned into foam cells (**Figures 2B-C**). Cholesterol efflux was also promoted by isolated HDL (**Figure 2D**). This effect was blunted after VSMC cholesterol-loading but enhanced following activation with an LXR agonist (**Figure 2D).**

**Figure 2:**
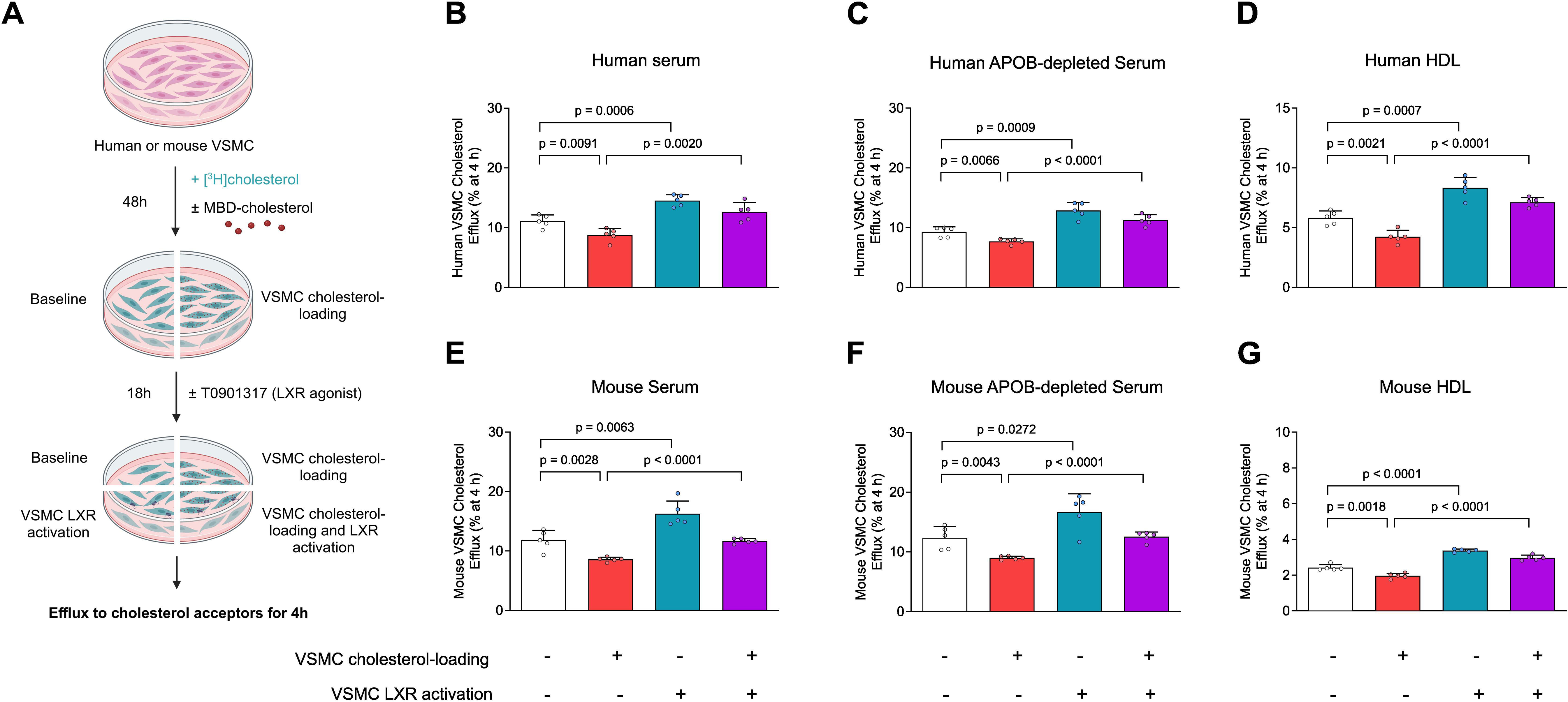
LXR Agonist Enhances Cholesterol Efflux in VSMCs, Including Those Exposed to MBD-Cholesterol, *In Vitro*. (A) Schematic representation of the VSMC cholesterol efflux assay. Primary VSMCs derived from human aortas and immortalized mouse aortic VSMCs were loaded with [1α,2α(n)-^3^H]cholesterol under baseline conditions or during exposure to 20 μg/mL MBD-cholesterol, representing the VSMC cholesterol-loading phase, for 48 hours. VSMCs were then treated with either the vehicle or 2 µmol/L of the LXR agonist T0901317 for 18 hours and subsequently exposed to various human and murine cholesterol acceptors for 4 hours. (B) Cholesterol efflux from human VSMCs to serum (2.5% v/v). (C) Cholesterol efflux from human VSMCs to APOB-depleted serum (2.5% v/v). (D) Cholesterol efflux from human VSMCs to HDL (25 µ/mL). (E) Cholesterol efflux from mouse VSMCs to serum (2.5% v/v). (F) Cholesterol efflux from mouse VSMCs to APOB-depleted serum (2.5% v/v). (G) Cholesterol efflux from mouse VSMCs to HDL (25 µg/mL). Data are presented as the mean ± SD from 5 independent experiments per group. Unpaired t-tests were used to compare cholesterol efflux between groups. APO: apolipoprotein, HDL: high-density lipoprotein, LXR: Liver X Receptor, MBD: methyl-β-cyclodextrin, VSMC: vascular smooth muscle cell.

These changes in VSMC cholesterol efflux followed a similar pattern when mouse aortic VSMCs were transitioned into foam cells and treated with the LXR agonist (**Figure 2E-G).** We further evaluated the expression of *ABCA1* in human VSMCs and in those exposed to MBD-cholesterol following treatment with an LXR agonist (**Figure S1**). *ABCA1* expression was upregulated during the transition into foam cells, and this upregulation was further enhanced by LXR agonist treatment (**Figure S1A**). These changes in *ABCA1* followed a similar pattern when mouse aortic VSMCs were loaded with MBD-cholesterol and treated with the LXR agonist (**Figure S1B**).

### LXR agonist treatment in mouse VSMCs is sufficient to substantially promote RCT to feces *in vivo*

To test whether activation of LXR in mouse VSMCs is sufficient to promote RCT in vivo, [^3^H]cholesterol-labeled mouse VSMCs that were loaded with MBD-cholesterol or not, were pretreated with either vehicle or the LXR agonist and then injected into wild-type male mice fed a Western-type diet for 4 weeks (see the experimental outline in **Figure 3A**). We assessed the VSMC-to-feces rate *in vivo* by evaluating [^3^H]cholesterol recovery in serum, HDL, and liver at 48h, along with the feces collected over 48h (**Figure 3A**). As expected, the injection of radiolabeled mouse VSMCs, whether loaded with MBD-cholesterol and pretreated with the LXR agonist or not, had no impact on the systemic serum lipoprotein profile or liver lipid levels in mice (**Table S1)**. However, the accumulation of VSMC-derived [^3^H]cholesterol in serum, HDL, and liver was reduced following the injection of radiolabeled mouse VSMCs loaded with MBD-cholesterol (**Figure 3B-D**). Additionally, VSMC cholesterol-loading markedly decreased the transfer of radiolabeled cholesterol to feces in mice (**Figure 3E**). Notably, the injection of [^3^H]cholesterol-labeled mouse VSMCs pretreated with the LXR agonist enhanced the transfer of radiolabeled cholesterol to serum, HDL, liver and feces compared to mice injected with untreated mouse VSMCs (**Figure 3B-E**). Similar changes were observed when VSMCs loaded with MBD-cholesterol and pre-treated with the LXR agonist were compared to untreated MBD-cholesterol-loaded VSMCs, except for [^3^H]cholesterol accumulation in the liver, which showed a notable trend toward an increase (**Figure 3B-E**).

**Figure 3:**
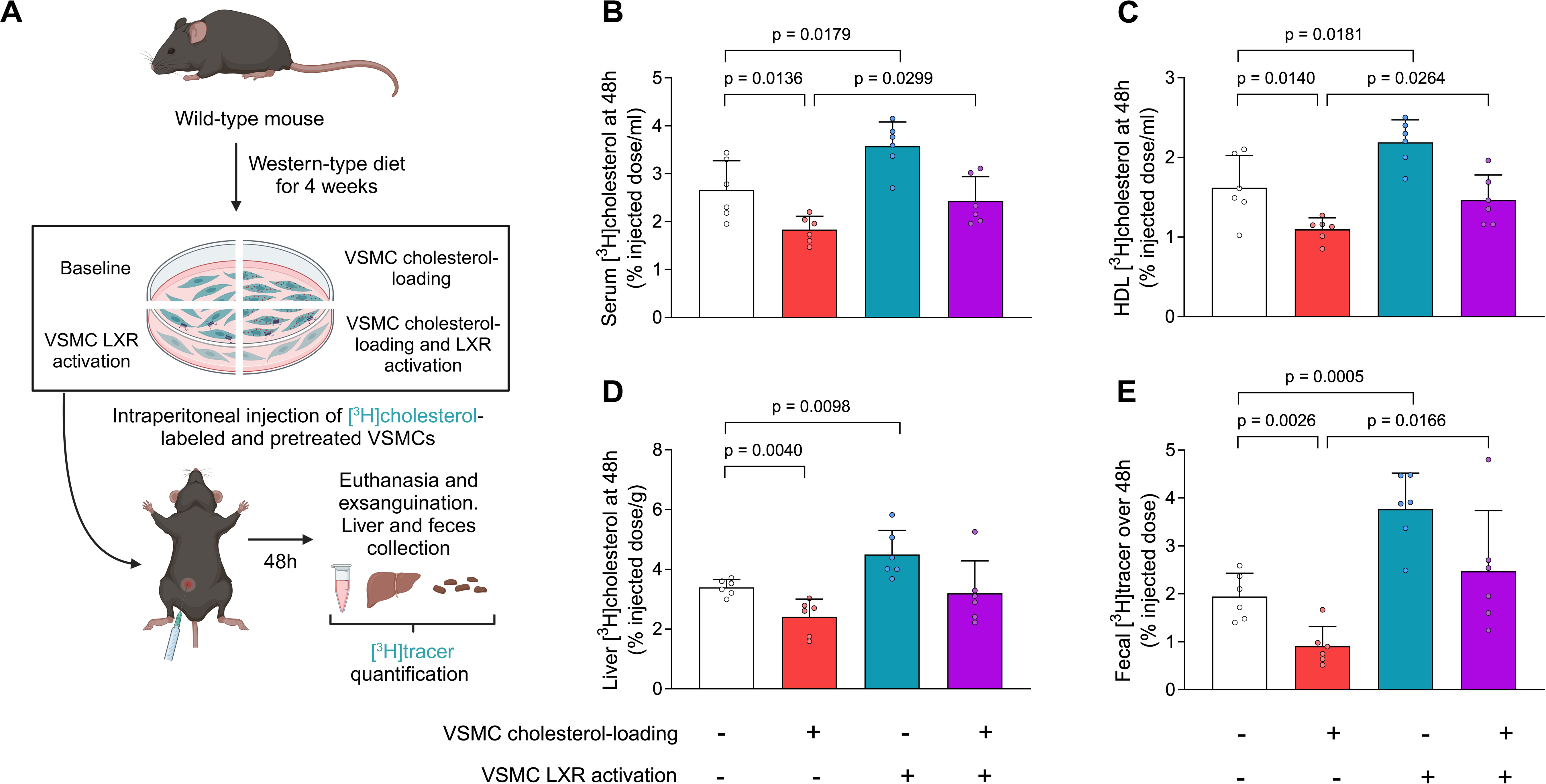
Selective LXR Agonist Treatment in Mouse VSMCs Promotes RCT to Feces *In Vivo*. (A) Schematic representation of the VSMC-to-feces RCT assay. Eight- to ten-week-old male mice were fed a Western-type diet for 4 weeks. Immortalized mouse aortic VSMCs were labelled with [1α,2α(n)-^3^H]cholesterol under baseline conditions or during exposure to 20 μg/mL MBD-cholesterol, representing the VSMC cholesterol-loading phase, for 48 hours. VSMCs were then treated with either the vehicle or 2 µmol/L of the LXR agonist T0901317 for 18 hours. Forty-eight hours before the end of the experiment, mice were randomized into four groups and intraperitoneally injected with the radiolabeled mouse VSMCs. The groups were as follows: 1. Mice injected with radiolabeled VSMCs under baseline conditions. 2. Mice injected with radiolabeled VSMCs under cholesterol-loading conditions. 3. Mice injected with radiolabeled VSMCs pre-treated with the LXR agonist. 4. Mice injected with radiolabeled VSMCs under cholesterol-loading conditions and pre-treated with the LXR agonist. Mice were housed in individual cages. At the end of the experiment, they were euthanized via cardiac puncture for exsanguination, and their livers and stool samples were collected for analysis. (B, C) [^3^H]cholesterol levels in serum and HDL at 48 hours post-injection. (D) [^3^H]cholesterol levels in the liver at 48 hours post-injection. (E) [^3^H]tracer in feces collected over 48 hours. Values are expressed as the mean ± SD of 6 animals per group. The amount of [^3^H]tracer is expressed as a fraction of the injected dose. Unpaired t-tests were performed to compare serum [^3^H]cholesterol, HDL [^3^H]cholesterol, liver [^3^H]cholesterol, and fecal [^3^H]tracer levels between groups. HDL: high-density lipoprotein, LXR: Liver X Receptor, MBD: methyl-β-cyclodextrin, RCT: reverse cholesterol transport, VSMC: vascular smooth muscle cell.

The LXR agonist also enhanced the overall transport of [^3^H]cholesterol from both mouse VSMCs and cholesterol-loaded mouse VSMCs to feces in wild-type female mice fed the Western-type diet for 4 weeks (**Figure S2**).

### Systemic LXR agonist treatment also promotes VSMC-to-feces RCT

To further evaluate the systemic effect of the LXR agonist, wild-type male mice were gavaged with the LXR agonist or vehicle for 10 days prior to the injection of mouse VSMCs or VSMCs loaded with MBD-cholesterol. Gavage was continued throughout the 48-hour experimental period (see the experimental outline in **Figure 4A**). At the systemic level, serum cholesterol and HDL cholesterol levels were increased in LXR agonist-treated mice (**Table S2**). LXR agonist treatment significantly increased liver triglyceride levels while reducing cholesterol levels (**Table S2)**. Notably, when [^3^H]cholesterol-labeled mouse VSMCs and those loaded with MBD-cholesterol were injected into mice treated with the LXR agonist, there was a higher accumulation of radiolabeled cholesterol in serum, HDL, and feces compared to mice treated with the vehicle solution (**Figure 4B, C, and E**). However, the accumulation of VSMC-derived radiolabeled cholesterol in the livers of mice was reduced following treatment with the LXR agonist (**Figure 4D**).

**Figure 4:**
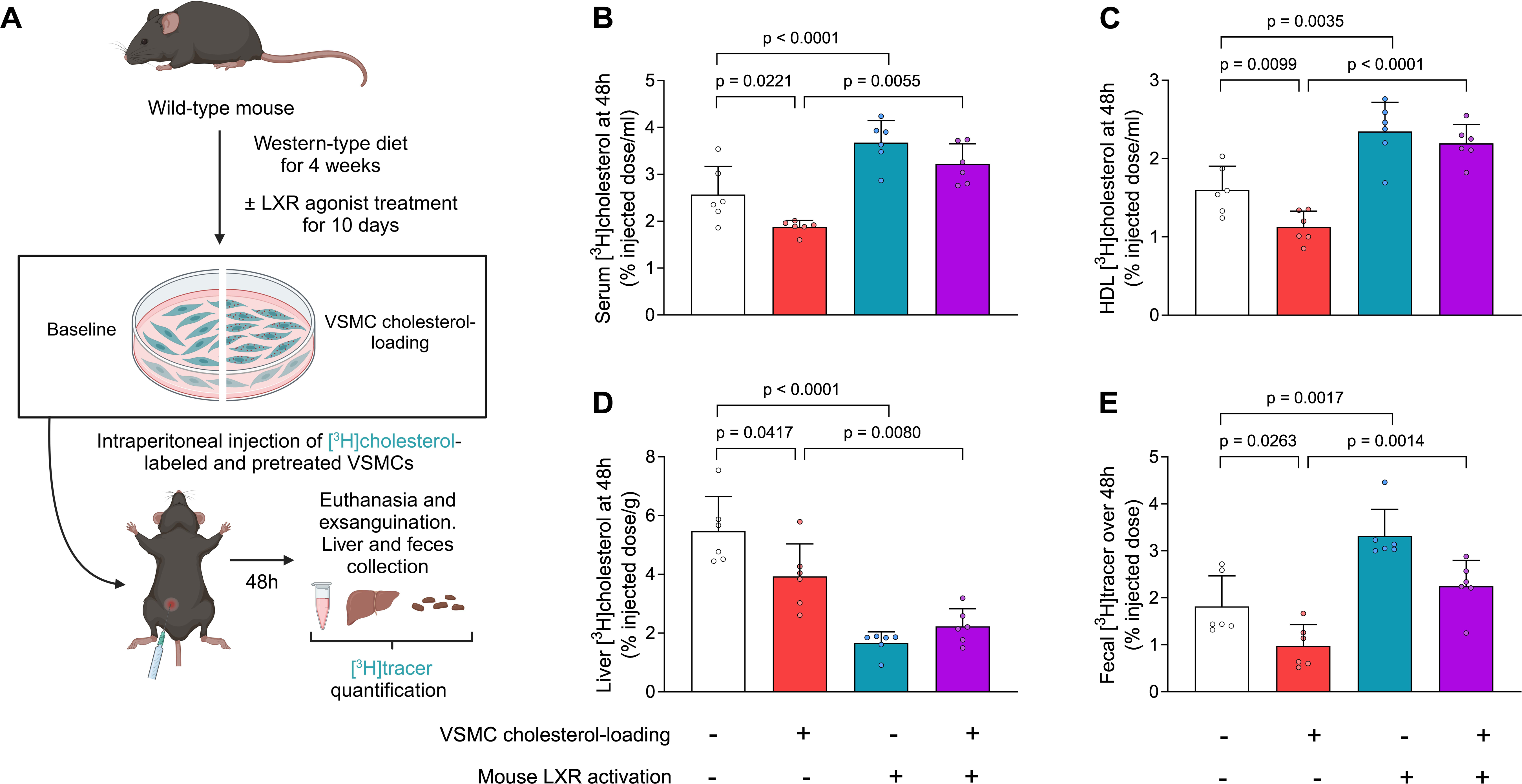
Systemic LXR Agonist Treatment Enhances RCT from VSMCs to Feces *In Vivo*. (A) Schematic representation of the VSMC-to-feces RCT assay in LXR-treated mice. Eight- to ten-week-old male mice were fed a Western-type diet for 4 weeks. Immortalized mouse aortic VSMCs were labelled with [1α,2α(n)-^3^H]cholesterol under baseline conditions or during exposure to 20 μg/mL MBD-cholesterol, representing the VSMC cholesterol-loading phase, for 48 hours. Mice were divided into two groups and treated with either a vehicle solution (1% carboxymethylcellulose) or the LXR agonist T0901317 (10 mg/kg in 1% carboxymethylcellulose) during the final 12 days of the diet, including the 48 hours of the RCT assay. Forty-eight hours before the end of the experiment, each group was further randomized into two additional groups and intraperitoneally injected with radiolabeled mouse VSMCs. The groups were as follows: 1. Mice treated with the vehicle solution and injected with radiolabeled VSMCs under baseline conditions. 2. Mice treated with the vehicle solution and injected with radiolabeled VSMCs under cholesterol-loading conditions. 3. Mice treated with the LXR agonist and injected with radiolabeled VSMCs under baseline conditions. 4. Mice treated with the LXR agonist and injected with radiolabeled VSMCs under cholesterol-loading conditions. Mice were housed in individual cages. At the end of the experiment, they were euthanized via cardiac puncture for exsanguination, and their livers and stool samples were collected for analysis. (B, C) [^3^H]cholesterol levels in serum and HDL at 48 hours post-injection. (D) [^3^H]cholesterol levels in the liver at 48 hours post-injection. (E) [^3^H]tracer in feces collected over 48 hours. Values are expressed as the mean ± SD of 6 animals per group. The amount of [^3^H]tracer is expressed as a fraction of the injected dose. Unpaired t-tests were performed to compare serum [^3^H]cholesterol, HDL [^3^H]cholesterol, liver [^3^H]cholesterol, and fecal [^3^H]tracer levels between groups. HDL: high-density lipoprotein, LXR: Liver X Receptor, MBD: methyl-β-cyclodextrin, RCT: reverse cholesterol transport, VSMC: vascular smooth muscle cell.

### ACAT inhibition normalizes the rate of RCT from VSMCs loaded with MBD-cholesterol to feces *in vivo*

To further investigate the mechanism by which MBD-cholesterol reduces the rate of VSMC-to-feces RCT, we evaluated the bioavailability of VSMC radiolabeled unesterified cholesterol and examined the ability of VSMCs, loaded with MBD-cholesterol and treated with an ACAT inhibitor, to mobilize and transport cholesterol to serum and feces (see the experimental outline in **Figure 5A**). Thin-layer chromatography analyses revealed that VSMCs retained over 94% of the radiolabeled cholesterol in its unesterified form, except for those VSMCs exposed to MBD-cholesterol, which reduced the bioavailability of unesterified radiolabeled cholesterol to 75% (**Figure 5B**). Notably, VSMC cholesterol-loading combined with ACAT inhibitor treatment fully restored the availability of unesterified radiolabeled cholesterol (**Figure 5B**). The ACAT inhibitor also completely normalized the ability of MBD-cholesterol-loaded VSMCs to induce cholesterol efflux to serum, an effect that was further accelerated by the LXR agonist (**Figure 5C**). When VSMCs loaded with MBD-cholesterol and treated with an ACAT inhibitor were injected into mice, the levels of radiolabeled cholesterol in serum and feces were similar to those of unloaded and untreated VSMCs (**Figure 5D and E**). Additionally, the LXR agonist further enhanced the overall transport of [^3^H]cholesterol from MBD-cholesterol-loaded mouse VSMCs treated with the ACAT inhibitor (**Figure 5D and E**).

**Figure 5:**
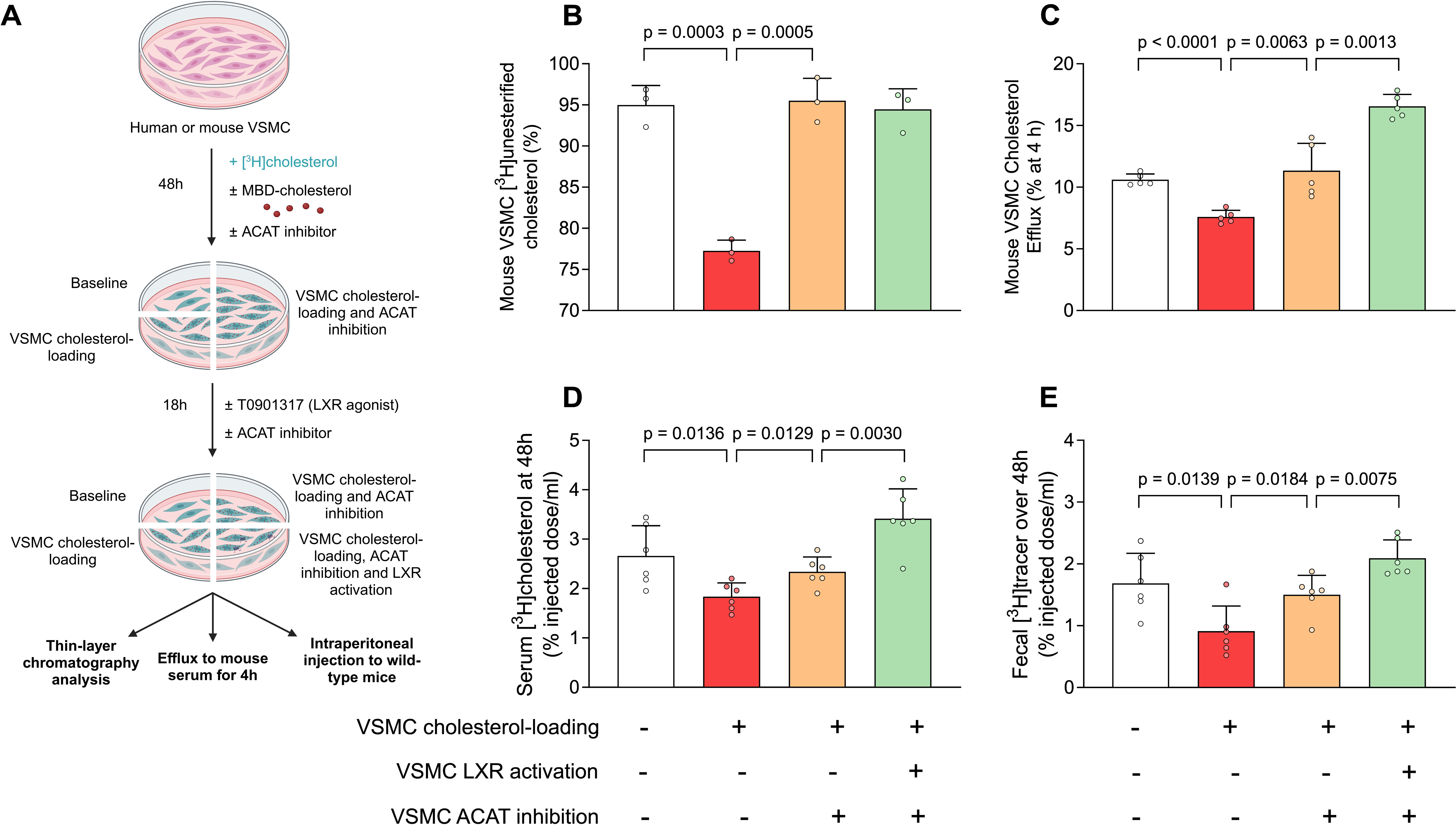
Selective ACAT Inhibition Restores VSMC Cholesterol Efflux and the Rate of RCT from MBD-Cholesterol-Loaded VSMCs to Feces. (A) Schematic representation of the VSMC experiments with the ACAT inhibitor. Immortalized mouse aortic VSMCs were loaded with [1α,2α(n)-³H]cholesterol under baseline conditions or exposed to 20 μg/mL MBD-cholesterol, representing the VSMC cholesterol-loading phase, for 48 hours. Simultaneously, VSMCs were treated with or without the ACAT inhibitor Sandoz 58-035 (10 µmol/L). Subsequently, cholesterol-loaded VSMCs were treated with either the vehicle or 2 µmol/L of the LXR agonist T0901317 for 18 hours, also in the presence of the ACAT inhibitor. Lipids were extracted from VSMCs, and thin-layer chromatography analyses were performed to measure the distribution of cellular [³H]unesterified and esterified cholesterol. VSMC cholesterol efflux and VSMC-to-feces RCT assays were conducted as described in Figure 2 and 3. The experimental groups were as follows: 1. Radiolabeled VSMCs under baseline conditions. 2. Radiolabeled VSMCs under cholesterol-loading conditions. 3. Radiolabeled VSMCs under cholesterol-loading conditions and pre-treated with the ACAT inhibitor. 4. Radiolabeled VSMCs under cholesterol-loading conditions and pre-treated with the ACAT inhibitor and the LXR agonist. (B) Percentage of cellular [³H]unesterified cholesterol relative to [³H]total cholesterol. Data are presented as the mean ± SD from 3 independent experiments per group. (C) Cholesterol efflux from human VSMCs to serum (2.5% v/v). Data are presented as the mean ± SD from 5 independent experiments per group. (D) VSMC-to-feces RCT assay: [³H]cholesterol levels in serum 48 hours post-injection. (E) VSMC-to-feces RCT assay: [³H]tracer in feces collected over 48 hours. (D, E): Data are presented as the mean ± SD from 6 mice per group. Unpaired t-tests were used to compare the percentage of cellular [³H]unesterified cholesterol and RCT metrics between groups. ACAT: acetyl-CoA acetyltransferase, LXR: Liver X Receptor, MBD: methyl-β-cyclodextrin, RCT: reverse cholesterol transport, VSMC: vascular smooth muscle cell.

## Discussion

The present study builds upon the growing recognition of VSMCs as a significant source of foam cells in atherosclerotic lesions, contributing to 30–70% of foam cell populations.^4,5,12,20–22^ While the activation of LXRs has been well-established in enhancing RCT from macrophages *in vivo*,^15,16,23^ its role in regulating the entire VSMC-specific RCT pathway remains unknown.

This study addresses this critical gap by evaluating the impact of LXR activation on cholesterol efflux and RCT from VSMCs, as well as in foam-like cells derived from VSMCs. Initially, we demonstrate that serum and HDL induce cholesterol efflux from both primary human and immortalized mouse aortic VSMCs at comparable levels, establishing cross-species consistency. Notably, cholesterol efflux capacity was significantly impaired upon MBD-cholesterol loading, which mimics key features of foam cell formation. This impairment highlights a substantial disruption in cholesterol handling in foam-like VSMCs.

Interestingly, we observed upregulation of *ABCA1* in both human primary and immortalized mouse VSMCs exposed to MBD-cholesterol. This observation contrasts with findings that *ABCA1* expression is significantly reduced in human intimal and rat intimal-like VSMCs.^4,9^ However, other studies have reported increased *Abca1* mRNA expression in cholesterol-loaded mouse VSMCs.^5,6,24^ This discrepancy suggests that cholesterol loading can stimulate *ABCA1* transcription in VSMCs under specific conditions, potentially as a compensatory mechanism to mitigate lipid accumulation. Nonetheless, despite this upregulation, the levels of *ABCA1* appear insufficient to prevent cholesterol accumulation in VSMCs under hypercholesterolemic conditions.

Treatment with the LXR agonist T090137 significantly enhanced cholesterol efflux from both human and mouse VSMCs. These findings suggest that LXR activation effectively upregulates key cholesterol transport pathways, potentially involving transporters such as ABCA1. Notably, although LXR-mediated enhancement was evident, cholesterol efflux from lipid-laden VSMCs remained less efficient compared to non-lipid-laden VSMCs. This observation underscores intrinsic limitations in the capacity of foam cells to mobilize cholesterol effectively. In contrast, a complete failure of LXR-regulated cholesterol efflux was observed in intimal-like VSMCs derived from the arteries of Wistar-Kyoto rats, potentially due to the absence of a critical cell surface or cytoskeletal component necessary for APOA1 binding to these VSMCs.^9^ These findings underscore potential species-specific differences in the regulation of cholesterol efflux in VSMCs, highlighting the need for further investigation to elucidate the regulatory factors underlying these disparities in cultured VSMCs.

Consistent with our *in vitro* findings, *in vivo* experiments demonstrated impaired RCT from radiolabeled MBD-cholesterol-loaded VSMCs. Specifically, reduced cholesterol transfer to serum, HDL, liver, and feces was observed compared to non-lipid-laden VSMCs, reflecting the physiological challenges associated with cholesterol removal from VSMC-derived foam cells. Notably, pre-treatment of VSMCs with the LXR agonist significantly increased radiolabeled cholesterol levels in serum, HDL, and fecal cholesterol excretion, indicating a marked improvement in RCT in mice. Furthermore, systemic LXR activation enhanced cholesterol transport from untreated VSMCs to serum and feces. However, this improvement was accompanied by a reduction in radiolabeled cholesterol accumulation in the liver. This reduction may be attributable to liver LXR activation promoting cholesterol biliary excretion, thereby decreasing hepatic cholesterol storage.^25^ It should be noted that systemic LXR activation is harmful as it promotes lipogenesis and liver lipid accumulation,^26^ increasing the risk of hepatic steatosis. Overall, our findings highlight selective LXR activation in VSMCs as a promising strategy to enhance cholesterol removal without adverse liver effects.

The full restoration of VSMC cholesterol efflux and RCT rates following pre-treatment with an ACAT inhibitor highlights the intricate interplay between intracellular cholesterol esterification and efflux pathways. Unlike the toxic effects of ACAT inhibitors on foam cell formation in mouse macrophages—where unesterified cholesterol accumulation induces cytotoxicity^27,28^—ACAT inhibition has been shown in numerous studies to reduce total cholesterol accumulation in human macrophages by enhancing unesterified cholesterol efflux.^29–31^ Notably, human and rat aortic VSMCs exhibit resistance to the cytotoxic effects of cholesterol accumulation and ACAT inhibition, which effectively reduces foam cell formation without inducing cell death.^32^ In our experiments, selective ACAT inhibition restored the bioavailability of radiolabeled unesterified cholesterol, while the combined application of an LXR agonist and an ACAT inhibitor synergistically enhanced cholesterol efflux and significantly promoted RCT. These findings underscore the limitations of RCT efficiency in heavily lipid-laden VSMCs and emphasize the potential of combination therapies to optimize cholesterol removal.

In this study, we utilized both human and mouse VSMCs to ensure translational relevance and integrated both *in vitro* and *in vivo* models to comprehensively assess cholesterol efflux and RCT. Nonetheless, certain limitations should be acknowledged. For practical reasons, the peritoneal cavity was employed as a surrogate for the arterial intimal environment. While small HDL particles can penetrate and interact with the peritoneal fluid^33^ the arterial intimal microenvironment in atherosclerotic lesions becomes more acidic and hypoxic. These conditions can significantly impact HDL remodeling and structural integrity.^34^ However, lymphatic vessels play a critical role in both the peritoneal cavity and the arterial intima by facilitating the transport of foam cell-derived cholesterol, incorporated into HDL particles, to the plasma during the RCT pathway.^35^

In conclusion, this study demonstrates that HDL-mediated cholesterol efflux is markedly impaired in VSMCs exposed to MBD-cholesterol, which have transitioned into foam cells. Pharmacological activation of LXRs significantly enhances RCT from VSMCs to feces *in vivo*, offering a potential therapeutic approach to reduce VSMC-derived foam cell formation. Furthermore, the combination of selective LXR agonists and ACAT inhibitors in VSMCs shows promise as a synergistic approach to restoring cholesterol homeostasis in lipid-laden VSMCs. These findings highlight the need for further research into these therapeutic interventions to address atherosclerosis and its associated complications.

## Supplemental Material

Figure S1-S2

Tables S1–S2

Data Set

## Abbreviations and acronyms

ABCA1: ATP-binding cassette subfamily A member 1
ACAT: acetyl-CoA acetyltransferase
APO: Apolipoprotein
DMEM: Dulbecco’s Modified Eagle’s Medium
GAPDH: Glyceraldehyde-3-phosphate Dehydrogenase
HDL: High-density lipoproteins
FBS: Fetal bovine serum
LXR: Liver X receptor
MBD: methyl-β-cyclodextrin
PBS: phosphate-buffered saline
RCT: Reverse cholesterol transport
TAGLN: Transgrelin
VSMC: vascular smooth muscle cell

